# Single cell derived organoids capture the self-renewing subpopulations of metastatic ovarian cancer

**DOI:** 10.1101/484121

**Authors:** Tania Velletri, Emanuele Carlo Villa, Michela Lupia, Pietro Lo Riso, Raffaele Luongo, Alejandro Lopez Tobon, Marco De Simone, Raoul J.P. Bonnal, Saverio Minucci, Stefano Piccolo, Nicoletta Colombo, Massimiliano Pagani, Ugo Cavallaro, Giuseppe Testa

**Author notes:** These authors contributed equally to the work (Tania Velletri, Emanuele Carlo Villa).

## Abstract

High Grade Serous Ovarian cancer (HGSOC) is a major unmet need in oncology, due to its precocious dissemination and the lack of meaningful human models for the investigation of disease pathogenesis in a patient-specific manner. To overcome this roadblock, we present a new method to isolate and grow single cells directly from patients’ ascites, establishing the conditions for propagating them as single-cell derived ovarian cancer organoids (scOCOs). By single cell RNA sequencing (scRNAseq) we define the cellular composition of metastatic ascites and trace its propagation in 2D and 3D culture paradigms, finding that scOCOs retain and amplify key subpopulations from the original patients’ samples and recapitulate features of the original metastasis that do not emerge from classical 2D culture, including retention of individual patients’ specificities. By enabling the enrichment of uniquely informative cell subpopulations from HGSOC metastasis and the clonal interrogation of their diversity at the functional and molecular level, this method transforms the prospects of precision oncology for ovarian cancer.

## Introduction

High Grade Serous Ovarian cancer (HGSOC) constitutes a major unmet need in oncology as one of the tumor types for which least progress has been made in the past decades. It has remained one of the most lethal gynecological cancers, due to the failure of surgery and chemotherapy at eradicating the disease and the ensuing nearly invariable recurrence^1,2^. This is, in turn, the result both of factors inherent to the biology of the disease and of technical limitations that have so far hampered its study.

The former relate, first of all, to the specific features of the anatomical district that enable very precocious dissemination through the abdomen, with metastatic ascities often concomitant with the first diagnosis. In addition, converging evidence points to the pharmacological resistance of the tumor propagating cells, varyingly referred to as cancer stem cells (CSCs) or cancer initiating cells (CICs), that can, thus, persist after chemotherapy and can often^3,4^ remain quiescent for months in the peritoneal cavity from which they fuel renewed and/or continuous growth^5,6^.

Among the technical hurdles that have hampered progress, the community has become increasingly aware of the inadequacy of available ovarian cancer cell lines to model physiopathologically relevant aspects of the disorder. Not only they do not allow, by definition, to correlate molecular aberrations to clinical histories^7^ and are thus of no use to advance the precision oncology agenda, but, at an even more basic level, most of the classically used lines feature genomic profiles that do not recapitulate the landscape of alterations observed in most primary tumor isolates^2,8,9^. The development of new methods to robustly capture, from the original lesions and in a patient-specific manner, the cell subpopulations that maintain tumor growth is, thus, a key priority in the field.

3D organoid cultures have recently emerged as a powerful modelling approach for a variety of disorders, offering the opportunity to recapitulate salient features of the original tissue or organ and propagate in vitro relevant subpopulations of cells representative of the original *in vivo* complexity^10^.

In cancer, organoids have been derived from several tumor types^11–15^, with colorectal cancer (CRC) organoids paving the way in demonstrating the transformative impact of such avatars in terms both of patient-specific modelling and of mechanistic insight into human tumor biology. The capacity to derive CRC organoids from individual tumor cells has been in this respect particularly salient, enabling to probe the mutational and functional diversification of individual tumor cells at unprecedented resolution^16^.

For HGSOC there are no organoid-based platforms that allow the prospective propagation and molecular characterization of individual tumor cells. Thus, we set out to establish a new method to isolate and grow them in 3D, taking advantage of the ability of HGSOC cancer initiating cells to grow in an anchorage-independent manner^17,18^ and harnessing this property for the establishment of clonal cultures in individual wells that could, thus, allow longitudinal tracing of their propagated features. Importantly, given the specific features of the disease recounted above, we reasoned that the method would be particularly relevant if applicable directly to metastatic ascitis, as a highly informative disease stage of relatively easy access for the streamlined translation of this method to the clinical setting. Finally, we used single cell RNA sequencing (scRNAseq) to define the cellular composition of HGSOC metastatic ascitis and trace its propagation in both 2D and 3D culture paradigms, finding that single cell-derived ovarian cancer organoids (scOCOs) recapitulate key features of the original metastasis that do not emerge from classical 2D culture, including retention of individual patients’ specificities. Thus, this method establishes the feasibility of enriching physiopathologically relevant cell subpopulations directly from HGSOC ascitis and clonally investigating their diversity at the functional and molecular level.

## Results

### Efficient derivation of ovarian cancer organoids (scOCOs) from single cells of metastatic ascites of HGSOC

Given the lack of physiopathologically meaningful, patient-matched models of HGSOC, we set out to establish a HGSOC modeling platform to allow (fig. 1): i) the streamlined and functionally based isolation of cancer propagating cells; ii) their growth into 3D organoids; iii) their serial propagation along with the computational reconstruction of its impact on modeling; and iv) the comparison of 2D and 3D cultures paradigms at single cell resolution to benchmark their ability to recapitulate HGSOC.

**Figure 1.**
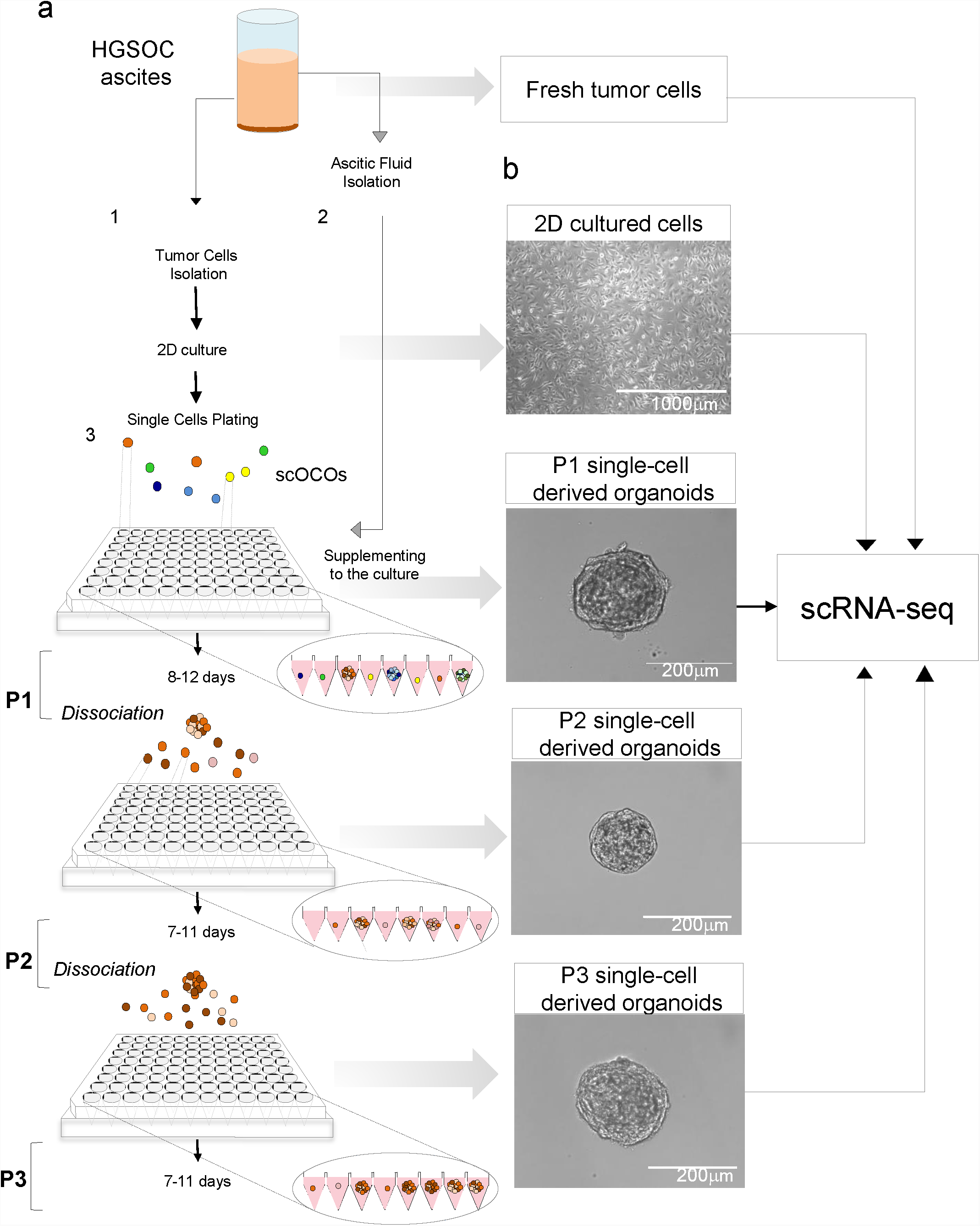
Generation of single cell organoids from HGSOC ascites. (a) Scheme illustrating the main steps of the method for generating single cell ovarian cancer organoids (scOCOS) from HGSOC ascites. Ascitic fluid is centrifuged; the cell pellet is processed for the removal of red cells and dissociated as single cell suspension for monolayer culture of tumor cells (step1); the remaining supernatant after the first centrifugation is processed in order to remove the residual fraction of cells and is used as supplement for growing and culturing scOCOs (step2). Tumor cells from monolayer culture are dissociated, resuspended in specific culture media for growing single cell from HGSOC ascites and plated by limiting dilution at the density of 1 cell per well in a low-adhesion 96 well plate V bottom (step 3). scOCOs at first passage (P1) are observed after 8-12 days in culture as tridimensional structure of about 200-220m of diameter and then propagated through dissociation of a single organoid in single cells. The single cells are resuspended in the described media and plated as previously indicated in order to obtain the next passages in culture (P2 and P3). The scheme illustrates that fresh tumor, monolayer culture of tumor cells, and scOCOs at different passages are then processed for scRNAseq to obtain their transcriptomic profile. (b) Bright field image analysis (x4magnification for 2D, scale bar 1000 □ m; x40 magnification for scOCOs P1, scale bar 400 □ m; x20 magnification for scOCOs P2 and P3, scale bar 200 □ m).

A hallmark of HGSOC is the ease with which it precociously metastasizes to the peritoneal cavity, which constitutes a key hindrance to its eradication. This dissemination is accompanied by the production of peritoneal fluid (ascites) whereas tumor cells can be present both as single cells and as floating aggregates. Given the impact of early and diffused metastasis on the poor management of HGSOC, we thus reasoned that ascites would be a particularly accessible and meaningful source of tumor cells for a translationally oriented HGSOC modeling platform.

To this end, we started by developing a method for isolating and culturing individual tumor cells from patients’ metastatic ascites (fig. 1, see Methods). Cancer Propagating Cells (CPCs), characterized by tumor-maintaining potential, self-renewal and anoikis-resistance, have been isolated from solid tumors and OC mostly by enrichment through tumor-sphere cultures, harnessing their ability to proliferate under non-adherent conditions^17,18^. For HGSOC, however, no method is yet available for the isolation and culture of individual tumor cells. Reasoning that ascites represents a particularly favorable niche for the growth of HGSOC CPCs, we supplemented the medium for growing primary cells with cells-free ascitic fluid at different ratios to define its baseline effect (supplementary fig. S1; methods). We observed a significant ascitic fluid-dependent increase in cell proliferation (supplementary fig. S2a-b) with the highest efficiency being 12.5% (Ascitic fluid: medium). We therefore selected this concentration as reference in setting up the optimal condition for the 3D culture of individual HGSOC cells. Specifically, HGSOC ascites from five previously untreated patients (Table 1-a) were processed in order to derive 2D cultures of tumor primary cells as described^19^ (fig. 2a). Cells from primary 2D cultures at the first *in vitro* passage were suspended in ovarian cancer stem cells medium with or without ascitic fluid (with the above-defined effective concentration) and plated by limiting dilution into 96 wells ultra-low attachment V-bottom plates at the density of one cell per well. We found that in such stringent non-adherent conditions no scOCOs are generated from single cell plated in medium alone (fig. 2b), while supplementation with cells-free ascitic fluid from patients enabled the proliferation of individual cells leading to the generation of scOCOs over a course of 8-12 days (fig 2c-d). These results indicate that cell-free ascitic fluid promotes cells proliferation in already in bidimensional cultures and is necessary to allow the 3D growth of individually seeded cancer propagating cells.

**Figure 2.**
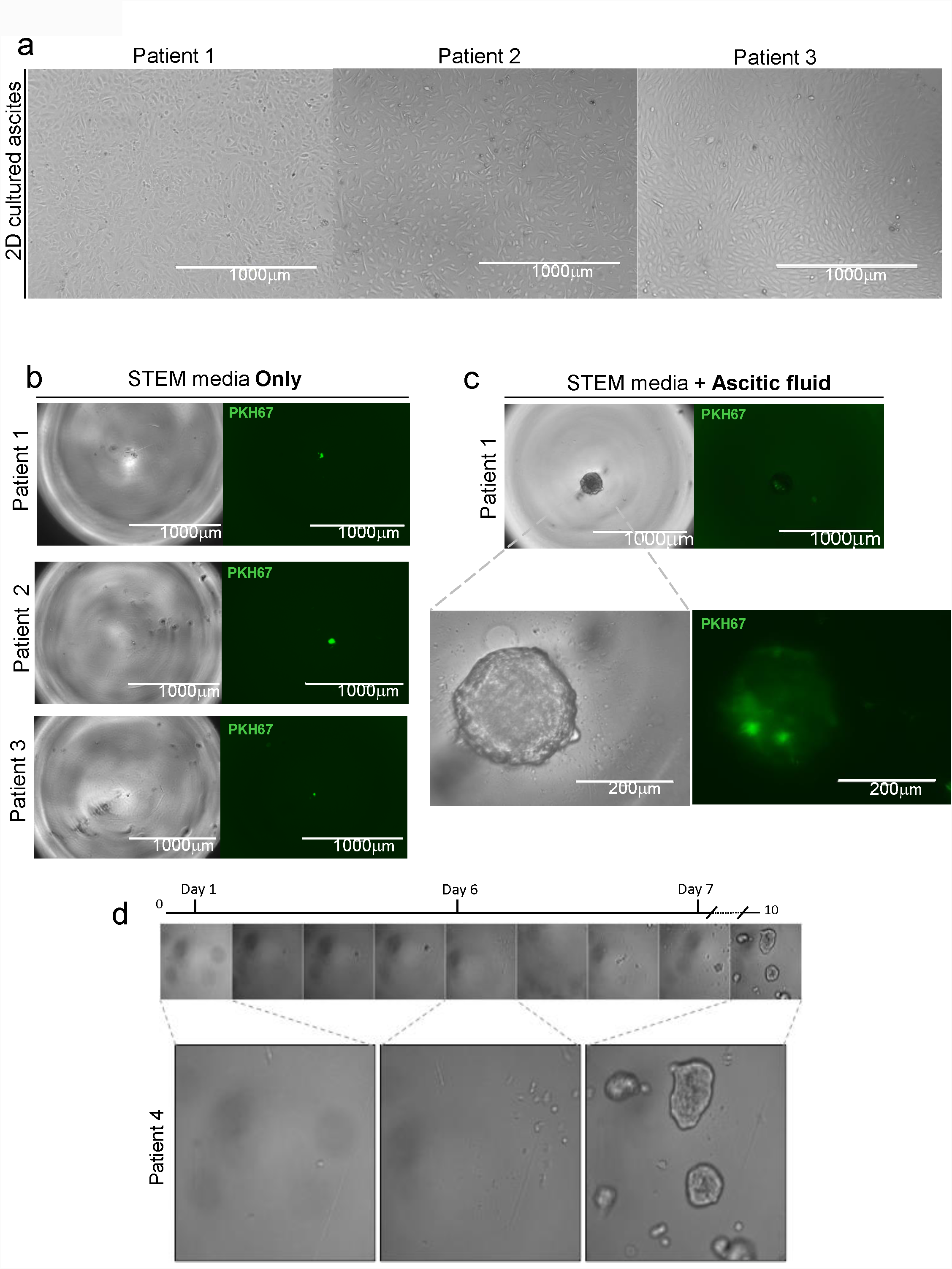
Metastatic ascitic fluid is required for cell proliferation and generation of single cell derived organoids. (a) Bright field images representative of the morphology of primary cultures derived from different HGSOC patients (x4 magnification for 2D monolayer culture of three different patients, scale bar 1000 □ m). **(b)** Bright field image analysis (x4 magnification for patient 1, patient 2 and patient 3, scale bar 1000 mm) of a single cell from 3 different patients plated in STEM media only and the respective fluorescent signal for PKH67+ (green). **(c)** Bright field image analysis (x4 magnification for patient 1, scale bar 1000 □ m; x20 magnification for patient 1, scale bar 200 □ m) of a single cell from patient 1 cultured in STEM media supplemented with ascitic fluid and fluorescent signal for PKH67. **(d)** Time-lapse of a single well followed during cell division. Day 0: only a single cell is present in the well. Day 2: first mitotic division. Day 4: the small organoid is formed. Day 5: the small organoids divide in single cells. Day 6: all the cells derived from the original organoids start to divide. Day 10: multiple monoclonal organoids are grown.

We then aimed to determine the efficiency of this process and the robustness of scOCOs across serial propagation. To ensure the monoclonal derivation of the organoid, primary cells were stained with the green fluorescent dye PKH67 for general cell membrane labeling methods, in order to be able to trace the single cell and monitor the formation of the organoid in a single well.

To this end, appropriate number of 2D primary cell cultures of patients with HGSOC (fig. 3a see methods) were labeled with the fluorescent dye and sorted by flow cytometry. The conditions for scaling down the number of cell doublets were optimized in order to obtain high purity (90-95%) of single labeled CPCs (see methods). Sorted cells were counted and plated by limiting dilution in a multiwell 96 well plate and supplemented with 12.5% of patients’ ascitic fluid.

**Figure 3.**
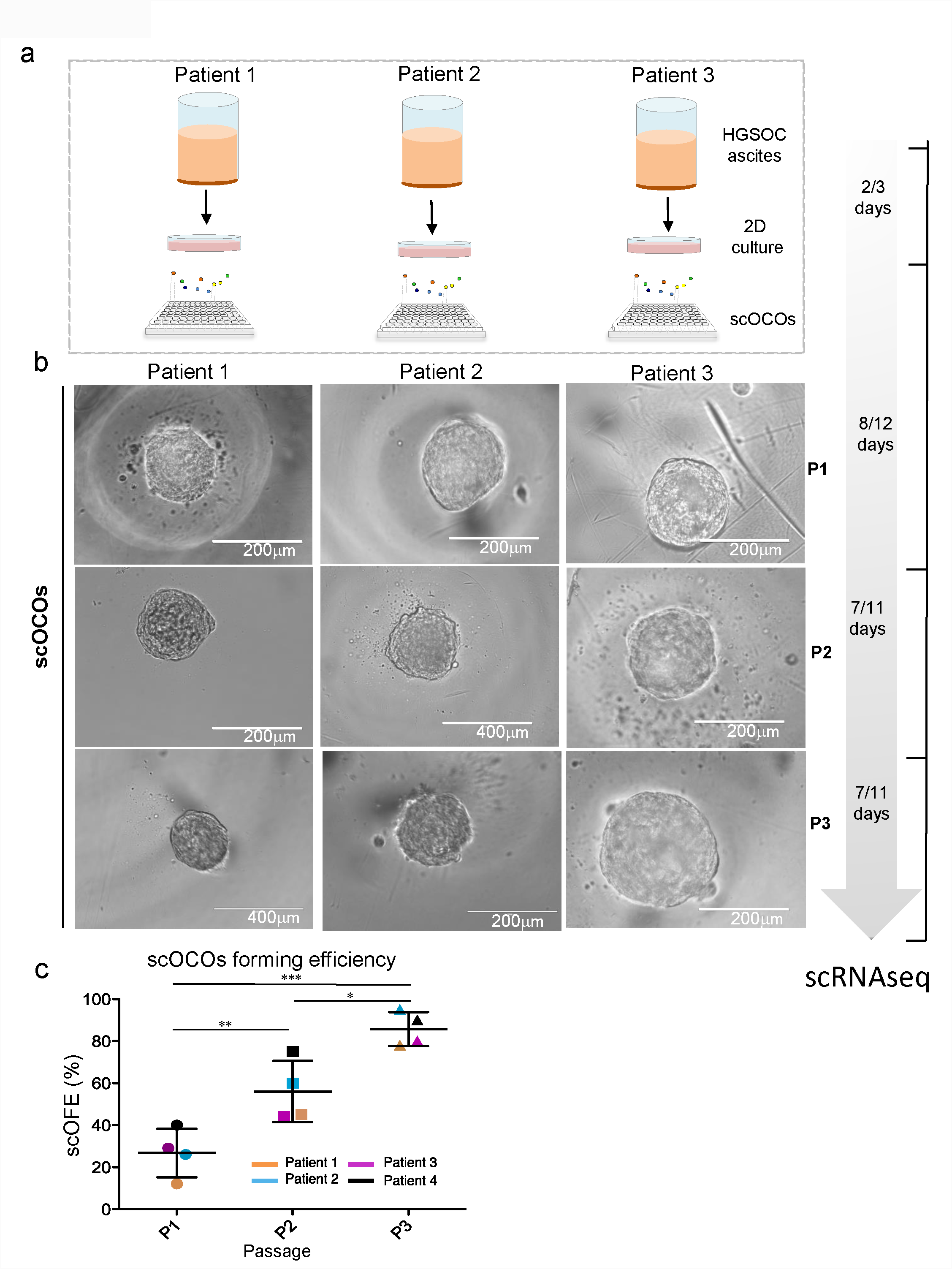
scOCOs forming efficiency increase during passages *in vitro*. (a) Scheme illustrating the main steps of the culturing method to derive scOCOCs with a dedicated timeline indicating the number of days required in culture for each stage.(b) Bright field image of representative images of scOCOs from three different patients at three different passages: P1,P2 and P3, (x20 magnification for scOCOs passage 1(P1) for patient 1, 2 and 3; forscOCOsP2 from patient 2 and 3; for scOCOsP3 from patient 3, scale bar 200 □ m); (x40 magnification for scOCOs passage 2 from patient 2 and for scOCOCs P3 from patient 1 and 2, scale bar 400 □ m). (c) Scatter plot showing the scOCOs forming efficiency (scOFE) as percentage for each patient and for each passage. Primary cultures derived from HGSOC ascites were grown under non-adherent conditions in 96 well plate V bottom in presence of the media supplemented with ascitic fluid to test their ability to generate monoclonal organoids. The experiment was performed on 4 independent samples. scOFE was calculated as the ratio between number of scOCOCs and the number of cells seeded. (d) Graph showing single cell organoid forming efficiency (scOFE) (mean + SEM) of scOCOs from different patients at passage 1, 2 and 3. P1 n=4, P2 n=4, P3 n=4. Unpaired t-test, (*) p<0.05; (**)p<0,01; (***) p<0.001). Table 2 is reporting the number of scOCOCs and the respective passage, for each patient, that were pooled to perform scRNAseq.

ScOCOs at passage 1 (P1) that reached a diameter of about 200-220 micron were dissociated into single cells and then plated again at the same conditions for propagation through passages 2 and 3 (P2 and P3 respectively) (fig. 3b). We observed that each subsequent passage dramatically increased scOCOs forming efficiency (fig. 3c), reaching 90-95% at passage 3, indicating a progressive enrichment for CPCs. Furthermore, by selecting and propagating, across three patients, two different P1 scOCOs and testing them separately, we also determined different efficiencies of propagation through P2 (fig. S3), thereby showing that different organoids from the same individual and at the same passage capture a gradient of propagation potentials. Finally, we tested the robustness of ascitic fluid supplementation by comparing patient-matched versus patient-unrelated experimental designs and confirmed their equivalence (fig. S4).

### Transcriptional dissection at single cell resolution of fresh metastatic tumors, 2D and 3D cultures

To define the cellular composition of scOCOs (3D) at single cell resolution and trace its dynamics over propagation *vis-à-vis* the original metastatic tumors and traditional 2D cultures, we performed single-cell RNA sequencing on a cohort of samples from five patients spanning the following conditions: freshly isolated HGSOC metastatic ascites, 2D culture of tumor primary cells and scOCOs at 2 different stages of propagation (P1 and P2) (fig. 4a).

**Figure 4.**
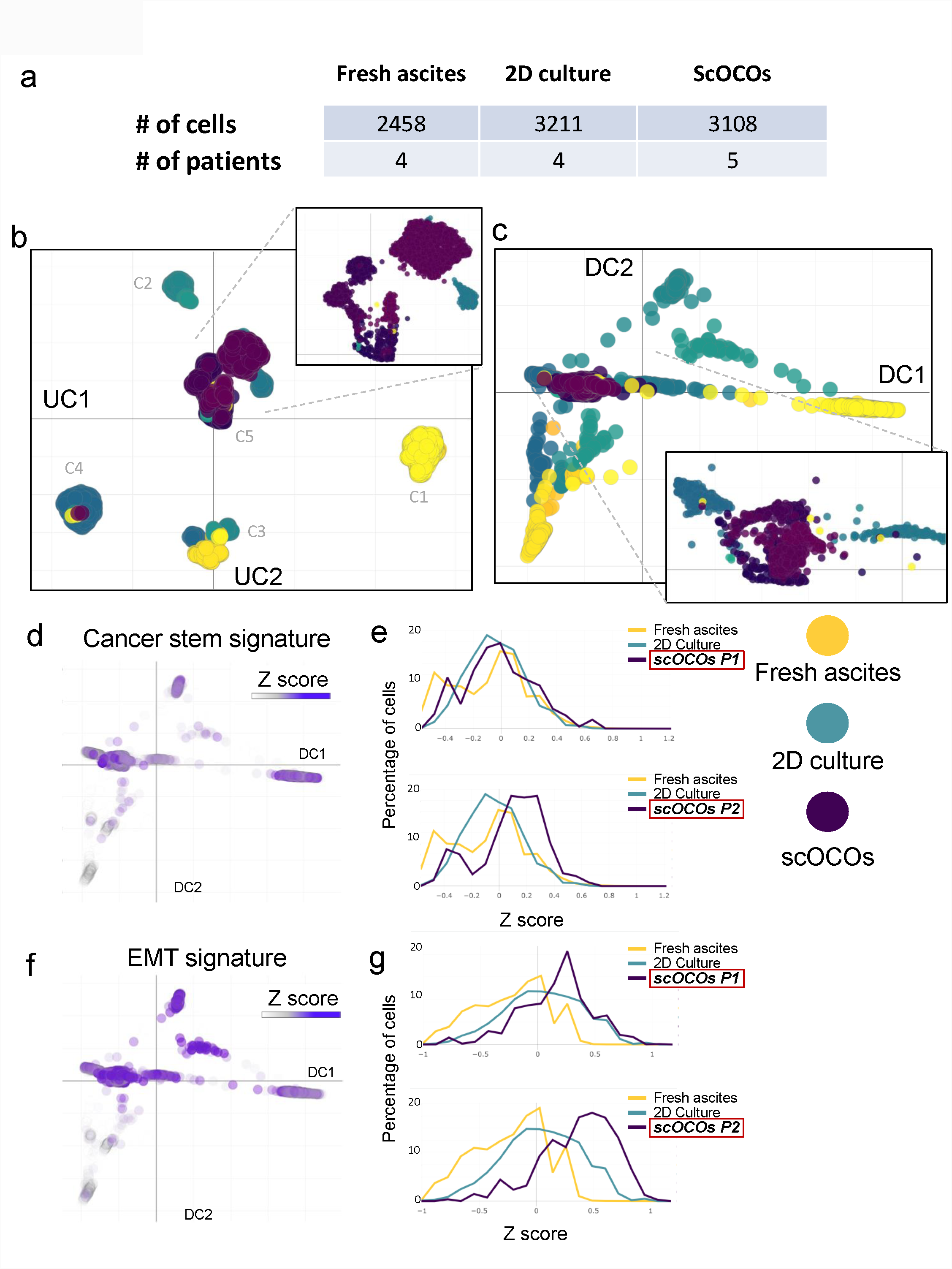
Single cell RNAseq of scOCOs reveals enrichment of CICs in scOCOs. (a) Number of cells and patients analyzed after filtering and quality control. (b)Umap of single cell transcriptomes from cells in (a), where each cell is represented by a point; yellow tones: fresh ascites; turquoise tones: 2D culture; purple tones: scOCOs; each color tone identifies different patients. Top right: the magnification of the central area of the Umap is enriched mainly for scOCOs cells but also for some ascites and 2D cultured cells. **(c)** Diffusion maps of single cell transcriptomes from cells in (a), where each cell is a point; yellow tones: fresh ascites; turquoise tones: 2D culture; purple tones: scOCOs; each color tone identifies different patients. Bottom right: magnification of the region where all the samples from different patients and conditions converge, enriched in scOCOs cells. (d,e,f,g) Enrichment analysis of Cancer stem and EMT signature. Left: diffusion maps of all the conditions and patients with a color scale defined by the z score of the respective signature; right: frequency plot showing the variation in the distribution of cells as a function of the z score of the indicated signature. The higher the percentage of cells with high z score, the more enriched the sample is for the indicated signature

In order to first obtain a global representation of the dataset we applied a non-linear dimensionality reduction visualization algorithm, Uniform Manifold Approximation and Projection (UMAP<u>^20^</u>), that has been shown to identify biologically meaningful cell clusters that retain consistency across a broad range of parameters variation, such as metric and number of neighbors. In addition, as an alternative to t-SNE, this approach also affords greater preservation of the global data structure^21^. This resulted in the identification of five different clusters of cells (fig. 4b), within which all cells belonging to the same cluster share specific characteristics. Cluster 1 (C1) is composed exclusively of freshly isolated tumor cells, cluster 2 (C2) of primary tumor cells cultured in 2D, cluster 3 (C3) contains both 2D cultured cells and freshly isolated cells, while cluster 4 (C4) is mainly composed by 2D cultured cells along with few fresh and scOCOs cells. Lastly, cluster 5 (C5) comprises cells belonging to all examined conditions, including the vast majority of scOCOs cells, indicating that scOCOs are homogeneous and that, importantly, they enrich for a specific subset of cells originally present in the fresh metastatic samples.

To examine the salient properties of our dataset in its entirety, we then applied diffusion map, a dimensionality reduction method that preserves the underlying structure of the original dataset, thus enabling a meaningful measure of the distances and trajectories intervening across any two given cells^22,23^ (fig. 4c). Diffusion map reveals a clear tri-partition, with the fresh HGSOC samples (in yellow) widely spread but clearly demarcated from the distributions of the 3D and 2D cultures (respectively in purple and turquoise, fig. 4c). This approach also confirmed a continuous relationship across samples represented by the overlap in the distribution of cells among the different conditions, consistent with the fact that freshly isolated tumor cells first undergo a single passage in 2D before being expanded in 3D. Importantly, this unsupervised approach also revealed that the two main components structuring the space of the diffusion map (DC1 and DC2) trace the specificity of the fresh samples from different patients, which however converge towards the region in which scOCOs cells are grouped, underscoring the consistency and homogeneity of features captured by the 3D culture system. Finally, both UMAP and diffusion map allow to draw two additional findings: i) a higher variability of 2D cultured primary cells compared to scOCOs; and ii) a high degree of consistency across scOCO passages, pointing to an enrichment in features that are specific of the 3D model.

### scOCOs capture specific features of metastatic HGSOC ascites

In order to investigate the cellular composition of scOCOs, we took advantage of validated cell-type specific gene markers to define transcriptional signatures to interrogate our dataset. To assess whether our 3D system is enriching for cancer initiating cells, we employed markers that have been widely used for their isolation in solid tumors and specifically in HGSOC^18^ (supplementary table 2). Moreover, considering that more than 90% of malignant ovarian tumors have an epithelial origin and that epithelial mesenchymal transition (EMT) is both a crucial factor for cancer progression and a prerequisite for metastatization^24,25^, we also investigated the expression of EMT-associated genes along with those defining the epithelial compartment *per se*^26,27^(supplementary table 2). The transcriptomic comparison between fresh metastatic samples, 2D and 3D cultures at different passages (fig.4a) revealed that scOCOs retain a consistent subpopulation of cancer initiating cells (fig. 4d), as measured by the Z score of the cancer stem signature reported in Supplementary table 2. Interestingly the number of cells with higher levels of expression of this signature is notably increased in P2 when compared to fresh tumor cells, 2D and scOCOs at P1, underscoring the progressive enrichment for stem cell features through propagation of the 3D model (fig. 4e). Furthermore, while we observed a higher expression of epithelial cells’ markers in fresh tumors when compared to 2D and scOCOs (supplementary fig. 5a-b), we found that the expression of EMT-related markers already increased in P1 scOCOCs, as compared to fresh cells and 2D cells, to reach an even higher level in P2 scOCOCs (fig. 4 f-g). Taken together, these results indicate that scOCOs, in contrast to 2D cultured cells, retain and progressively enrich for features of the fresh tumor related to cancer stemness and to the EMT process.

Finally, we sought to determine whether scOCOs could retain also patient-specific features, despite the overall convergence observed through diffusion map (fig. 4c). To this end, we performed a differential expression analysis in the whole dataset to identify genes that are differentially expressed in at least one of the patients irrespective of experimental condition ^28^. By this approach we identified 659 differentially expressed genes (DEGs), a subset of which, comprising 97 genes (15%), was retained between fresh tumor cells and scOCOs (supplementary fig. 6a). This suggests that scOCOs, despite being less variable than fresh and 2D cultures, retain a consistent portion of the gene expression profile characterizing each individual patient’s tumor. To confirm this result with an unsupervised approach, the same dataset was analyzed by weighted gene co-expression network analysis (WGCNA), selecting the gene modules most strongly associated to individual patients, regardless of condition. This led to the identification of 3 modules (supplementary fig. S6b, blue, green and greenyellow) which were then visualized by UMAP. As shown in fig S6c, UMAP clearly identifies patient-defining clusters that, importantly, comprise cells across all three conditions (fresh, 2D and 3D), thereby confirming that scOCOs are able to retain and propagate a salient portion of patient-specific tumor-associated gene expression.

### scOCOs highlight features related to HGSOC ascites that do not emerge from 2D culture

In order to assess whether scOCOs constitute a relevant and possibly superior alternative to classical 2D cultures, it was essential to verify which characteristics of the original metastatic samples were maintained throughout organoids’ propagation vis-à-vis the 2D counterpart. To test this, we used WGCNA to identify gene modules (Eigengenes, ie. genes whose change in expression relative to each other is consistently retained across individual cells) that correlate between conditions (fig. 5a), focusing on the features that are maintained between fresh ascites and 2D culture, and between fresh ascites and scOCOs. Among the identified modules none showed a strong correlation between 2D and scOCOs, whereas several were correlated between the fresh metastatic samples and, respectively, either the 2D or the 3D culture paradigms. In order to probe such similarities at the level of biological pathways, we applied gene ontology enrichment analysis on gene modules of either correlation.

**Figure 5.**
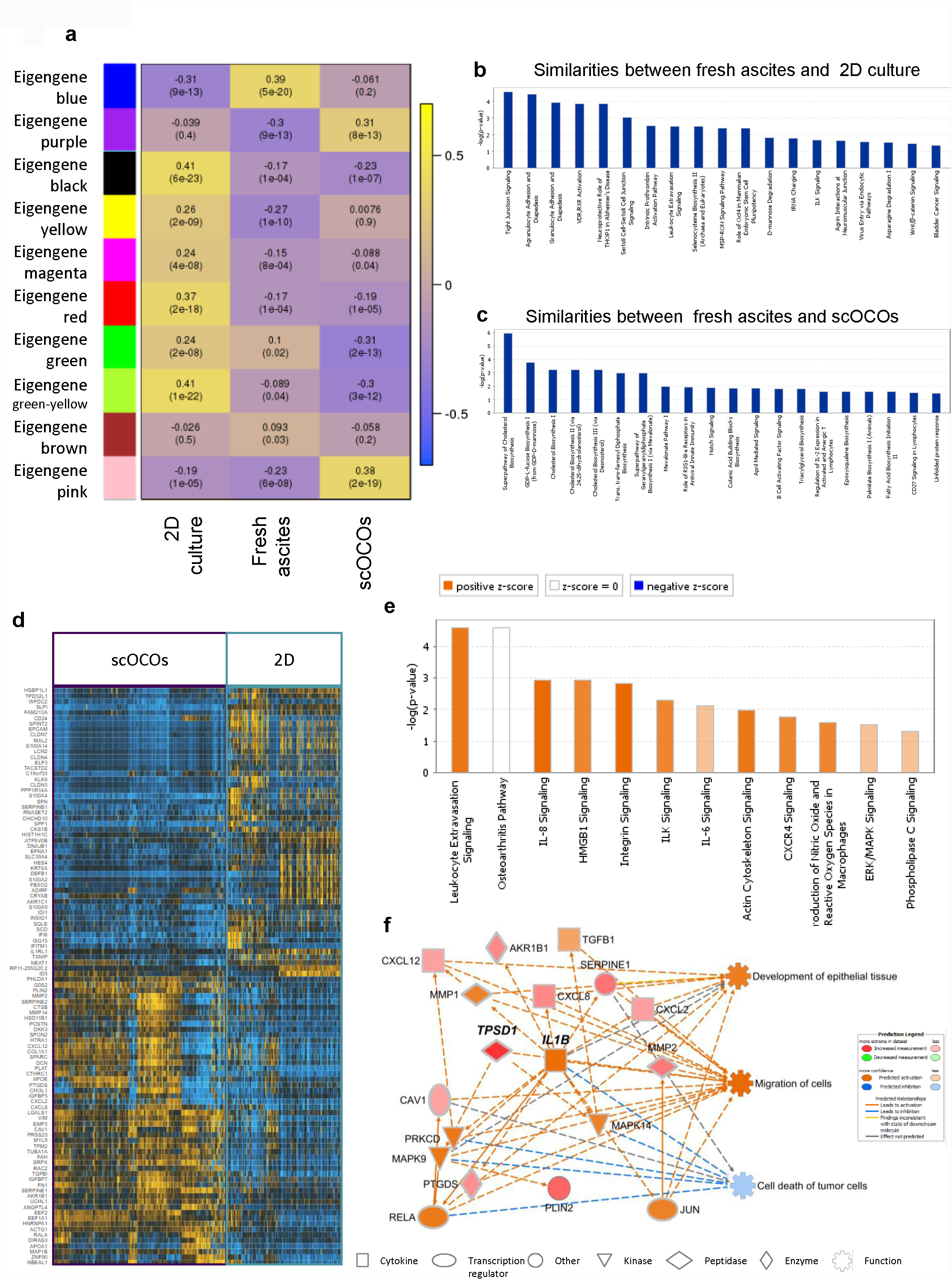
ScOCOs capture relevant features of the fresh tissue while preserving hallmarks of HGSOC ascites that do not emerge from 2D culture. (a) WGCNA labeled heatmap of whole transcriptome. Color maps define correlations between groups of genes and conditions; (b) pathway analysis on the groups of genes that correlate among fresh ascites and 2D culture; (c) pathway analysis on the groups of genes that correlate among fresh ascites and scOCOs; (d)heatmap of differentially expressed genes between scOCOs and 2D culture; (e) pathway analysis performed on DEGs in (d): color scale define the predicted activation/deactivation of the pathway, orange identifies the pathways activated in scOCOs; (f) IPA causal network derived from DEGs in (d): color scale define the expression of downstream regulated genes and their associated function, the orange tone identifies the genes and functions activated in scOCOs.

The analysis on the gene modules whose transcriptional behavior is shared between fresh ascites and 2D cultures revealed an enrichment for pathways related to signal transduction that regulate cell proliferation and gene expression^29^, diapedesis^30^, activation of vitamin D receptor pathway (VDR/RXR)^31,32^, as well as activity of integrin-linked kinase ILK^33^ all of which have been involved in regulation of tumor growth and also specifically ovarian cancer, as well as octamer binding transcription factor 4 (Oct-4) regulating stem cell self-renewal and pluripotency with emerging roles in regulating tumor initiating cells^34,35^ and Wnt beta-catenin signaling pathway (Wnt/□) involved in stem cell regeneration and organogenesis^36^ (fig.5b). In contrast, an ontology analysis of the genes defining the similarity between fresh ascites and scOCOs showed an enrichment in pathways related to the biosynthesis of cholesterol and triacyl glycerol biosynthesis. This is consistent with previous observations showing that abnormal expression levels and mutations of genes involved in the cholesterol homeostasis and lipid metabolism are related to cancer^37,38^ and OC^39,40^. Likewise, Notch signaling that regulates cell proliferation, stem cell maintenance and that plays a critical role in the cross talk between angiogenesis and CICs self renewal was also an enriched pathway defining the similarity between fresh and scOCOs cells^41^ (fig. 5c). Thus, our weighted gene co-expression network analysis (WGCNA) across conditions indicates that while 2D culture recapitulates well known biological pathways implicated in cell proliferation and tumor growth, scOCOs bring into relief additional features especially related to the metabolic and signaling state of the fresh metastatic samples that had so far resisted *in vitro* tractability.

Finally, to identify the specific differences between scOCOs and 2D cultured cells, we performed differential expression analysis between these two categories, identifying 104 DEGs common across all patients (fig. 5d). Functional analysis of this set of genes by Ingenuity Pathway Analysis (IPA)^42^ revealed an upregulation of pathways related to interleukin 8 (IL8)^43^ and integrin signaling in scOCOs, both correlating with tumor growth and progression (fig. 5e). Next, to investigate the upstream biological causes and predicted downstream effects of such differentially regulated circuits, we applied IPA causal network analysis (fig. 5f) and uncovered, in scOCOCs, the following two key insights: i) a downregulation of processes related to tumor cell death mediated through the action of v-rel avian reticuloendotheliosis viral oncogene homolog A also known as p65 (RELA), mitogen activated protein kinase 9 (MAPK9), mitogen activated protein kinase 14 (MAPK14), protein kinase C delta type (PRKCD) and interleukine 1 beta (IL1B); and ii) an up-regulation of the functions related to the development of epithelial tissue and cell migration, mainly mediated by IL1B, C-X-C motif chemokine 12 (CXCL12), C-X-C motif chemokine 8 (CXCL8) and tumor growth factor beta 1 (TGFB1). We can thus conclude that scOCOs retain and propagate functionally relevant features of fresh metastatic samples, including patient-specific ones, that do not emerge from current 2D culture systems.

## Discussion

We present here a new method for the derivation of 3D organoids from individual cells of HGSOC metastatic ascites. The method was designed to capture, from the diversity of cell subpopulations that characterize metastatic samples, the range of individual cells that are able to propagate tumor growth in 3D, thereby enabling the functional and molecular interrogation of metastatic HGSOC cell heterogeneity.

Our method presents the following key innovations.

First, we found that filtered ascitic fluid supplementation is strictly required to enable the growth of individual cells into scOCOs and their serial propagation. Importantly, this property holds across patients, pointing to the functional relevance of components shared among metastatic ascitis samples. Thus, while the identification of the specific molecule(s) mediating this effect will be an active area of mechanistic investigation, the method can be immediately implemented harnessing the generalized effect of ascitic fluid batches in streamlined, non patient-matched settings.

Second, the molecular characterization of scOCOs compared to fresh HGSOC ascites and 2D cultures through scRNAseq reveals that scOCOs retain and amplify, better than 2D cultures, specific cell subpopulations from the metastatic sample and that these reveal distinct and thus far undetected functional specificities (fig. 5a). Importantly, scOCOCs also retain a significant portion of the inter-patient molecular diversity detected in the fresh metastasis (fig. S6), a property that makes them relevant as sensitive, patient-matched avatars to advance precision oncology in the HGSOC field, both in terms of prognostic markers and druggable vulnerabilities.

Third, the very design of the scOCO derivation set up provides clear edges over current approaches that have been interrogating ovarian cancer stemness relying on the generation of spheroids cultured in bulk^18^. A specific hallmark of OC is indeed the intraperitoneal metastatic route that cancer cells tread through the intra-abdominal fluid-filled space where they can survive either as single cells or as multi-cellular aggregates (MCAs)^44^. That such individual floating cells can persist and fuel metastatic growth and/or relapse underscores the need to uncover their properties at clonal resolution. In this respect, the scalability of our method affords particular advantages, being uniquely suitable for high throughput plating, using microwells or micropillars^45,46^ that allow to seed one cell at the time, thus optimizing the sample yield per experiment and thereby streamlining patient-specific drug screenings. scOCOs could thus be automatically seeded in well-dense plates and observed after controlled stimulation by drugs or molecular interference, isolating the best and worst responder for further multi-omics investigation. Such a highly scalable approach can generate large amounts of scOCOs from any patient, thus increasing statistical power for high-definition intra-patients studies.

Finally, the observation that scOCOs capture inter-patients tumor heterogeneity lays down the foundations, in HGSOC, for experimental pipelines aimed first at defining and then functionally probing the specificities of each patient’s tumor through robustly propagated *in vitro* avatars. Together, our results demonstrate the power of scOCOs in furthering the mechanistic dissection of metastatic HGSOC by aligning clinical reasoning to physiopathologically meaningful, experimentally tractable patients’ models.

## Online Methods

### Samples

Ascites samples were obtained upon informed consent from patients undergoing for surgery treatment for primary, not recurrent HGSOC, at the Gynecology Division of the European Institute of Oncology, IEO, Milan, Italy. Table 1-a contains the list of samples related to the patients diagnosis.

#### Ethical approval

The study was conducted upon approval of the Ethics Committee of the European Institute of Oncology (Milan) following its standard operating procedures (“presa d’atto” from 24/7/2017). Fresh tissue samples were obtained upon informed consent from patients undergoing surgery at the Gynecology Division of the European Institute of Oncology. Only tissue samples from patients who have given informed consent to i) the collection of samples for research purposes and their storage into the Biobank of the European Institute of Oncology and ii) the transfer of samples to other research institutions for cancer research purposes have been used in this project. Collected personal data have been pseudonymized, and have been stored and processed in compliance with the applicable data protection legislation, D.Lgs 196/2003 and, since 25 May 2018, Regulation (EU) 2016/679 (General Data Protection Regulation).

### Primary tumor cell culture

We isolated epithelial ovarian cancer (EOC) cells from metastatic peritoneal ascites of five patients according an already established protocol^19^. To derive primary tumor cells, obtained ascites were transferred in to polypropylene 750 ml Bio-bottle (Thermo-scientific cat.no. 75003699) centrifuged at 300 x g for 5 min, the supernatant was harvested for downstream processing, while the cells pellet was resuspended with ACK lysing buffer (Lonza, cat.no. 10-548E) and incubated for 5 minutes at RT to lysate red cells. Tumor-derived cells were cultured on collagen-I-coated cell culture flasks 75cm^2^ Corning BioCoat, cat.no. 354485) in CO^2^ incubator at 37 °C. All the primary cells in this work were used at passage 1 in culture.

### Ascites processing

According to the volume, the supernatant of the ascites obtained following the first centrifugation for the derivation of primary cells is transferred in 50 ml polypropylene tubes and centrifuged at (1900 x g) 3000 rpm for 30 minutes at RT in order to remove all residual cells^47^. The supernatant is collected and transferred into a disposable sterile filter system with bottle of 500ml and or 1000ml volume with a PES membrane of pore size of 0.22 micron for vacuum filtration (Sartorius stedim, Sartolab BT filter system cat.no.180C2, 180C3) while the residual pellet is discarded. The filtrate is directly used for experiments, or collected as working aliquotes in 15 ml tubes and stored at −80°C. After thawing, the filtrate is moved into a 50 ml tubes and filtrated again with a vacuum driven sterile filter of 50 ml volume (Millipore, Steriflip cat. no. SCGP00525) through a suction canister soft liners (Medline, MED-SOFT disposable pre-gelified liner).

### Generation of single cell derived organoids from HGSOC ascites

Primary tumor cells at passage 1 were dissociated through trypsin/EDTA (Lonza cat. no. BE17-161F), centrifuged at 300 x g for 5 min, the supernatant was removed and the cell pellet was washed twice with pre-warmed Dulbecco-s Phosphate Buffered Saline (D-PBS 1X) and incubated with 0.25% trypsin/EDTA solution at 37 °C until cells are detached from the bottom of the flask. Cells were collected in D-PBS, pelleted at 300 x g for 5 minutes at RT, suspended in 500 □ □l of pre-warmed stem media (MEBM) and counted both with TC- 20 automated cell counter (Bio-Rad) and counting chamber (Biosigma-Fast Read 102). The cells were centrifuged at 300 x g for 5 minutes, and the pellet was resuspended in pre warmed serum-free MEBM supplemented with 100U/ml penicillin, 100ug/ml streptomycin, 2mML-glutamine, 5ug/ml insulin, 0.5ug/ml hydrocortisone, 1U/ml heparin,2% B27, 20ug/ml epidermal growth factor, 20ug/ml fibroblast growth factor and diluted by serial limiting dilution at the density of 1 cell for well in the media supplemented with 12.5% of ascitic fluid for plating into low cell adhesion 96 well plates (Sumitomo Bakelite cat. no. MS- 9096V 96 well) with a final volume of 200 □ l for well.

Cells were monitored daily by microscope visualization, until organoids formation at passage 1. The time required for the growth of scOCOs is sample dependent from 7 to 12 days. No changing media is required, however if liquid evaporation occurred we supplemented 25 □ l-50 □ l of fresh media plus ascetic fluid. Percentage of single cell derived organoid forming efficiency (scOFE) was calculated for every passage as the ratio between the total number of organoids generated in a 96 well plate, and the number of cells seeded and expressed as percentage.

### Propagation of scOCOs in vitro

For propagation at P2 or P3, a single organoid at day 8-11 was moved from 96-well V-bottom ultra-low attachment plates to 48-well ultra-low attachment plates (Corning), and incubated with 400 □l of pre-warmed 0.25% trypsin/EDTA solution at 37 °C for 20-30 minutes with gently pipetting for 20 times every 8 minutes and visualized by microscope for the stadium of disaggregation. To avoid loose of cells the tip was always rinsed with medium before pipetting. Cells were harvested, washed once in MEBM supplemented with ascitic fluid and centrifuged at 300 x g for 5 min. Supernatant was discarded and the pellet of cells was re-seeded in 96 well plate as previously described in order to obtain passage 2, P2, which requires a culturing time of about 7-9 days. The same procedure was applied to generate scOCOs at passage 3, and organoid formation was observed also between 7- 9 days. Percentage of organoid forming efficiency was calculated accordingly.

### PKH67 staining

Epithelial adherent cells were dissociated as single cells, harvested, washed with MEBM media (Lonza cat. no. CC-3151) without serum centrifuged at 300 x g for 5 min. Cells were counted through with TC-20 automated cell counter (Bio-Rad). Cells were resuspended in a final working volume of 2ml following the procedures for general cell membrane labeling for PKH67 (Sigma-Aldrich cat. no. PKH67GL). We scaled-down the number of cells necessary for the staining in the final working volume of 2ml to 100.000 cells; EDTA 0.01% was added to the working. Cells were suspended in 2 ml of MEBM plus 0.01% EDTA.

### Imaging

Image were taken with EVOS Cell Imaging Systems, and images taken in fig2d were acquired in Leica SP5 confocal microscope with 10x objective. Image were taken every day for the first 9 days directly from the 96 wells. During the acquisition’s interval the samples were store in the incubator. Image were reconstructed by a custom script using python and OpenCV.

### FACS analysis

The percentage of positive stained PKH67 cells was measured by BD Influx Sorter (BD Biosciences). Positive stained cells were recovered after sorting in falcon round bottom polystyrene tubes (STEMCELL technologies cat.no. 38007) containing 5 ml MEBM plus 0.1% FBS.

### Single cell preparation, cDNAsynthesis generation of single cell GEM and libraries construction

Metastatic ascites, 2D primary cells and single cell derived organoids were dissociated in order to obtain single cells suspension. scOCOs were collected at day 10, and 20-25 organoids for pssage and for patient were dissociated by incubation with a solution 0.25% of trypsin/EDTA at 37 °C for 20 minutes. Cells in suspension were washed with D-PBS 1X and centrifuged at 300 x g for 5 min. The cell suspension was passed once through cell strainer (Bel-Art; Flowmi cell strainer for 1000 microliter pipette tips, cat. no. H13680-0040) to remove cellular debris and clumps and was resuspended with wide bored tip in D-PBS 1X supplemented with 0.04% bovine serum albumin BSA, Sigma. The cell concentration was determined through TC-20 automated cell counter (Bio-Rad). Droplet-based single cell partitioning to generate single cell gel beads in emulsion (GEMs) was obtained by loading the appropriate cell dilution onto 10x Genomics Single Cell 3’Chips mixed with reverse transcription mix using the Chromium Single-cell 3’ reagent kit protocol V2 (10x Genomics; Pleasanton, CA), according the manufacturer’s protocol. The gel beads are coated with unique primers bearing 10× cell barcodes, unique molecular identifiers (UMI) and poly(dT) sequences. The single-cell suspension at a density of 1000 cells/μl was mixed with RT-PCR master mix and loaded with Single-Cell 3′ gel beads and partitioning oil into a single-cell 3′ Chip. The chip was then loaded onto a Chromium instrument for single-cell GEM generation within which RNA transcripts from single cells are reverse-transcribed. Barcoded full-length cDNA are generated using Clontech SMART technology. cDNA molecules from one sample were pooled and preamplified. The amplified cDNAs were fragmented. Final libraries were incorporated with adapters and sample indices compatible with Illumina sequencing, and quantified by real-time quantitative PCR (calibration with an in-house control sequencing library). The size profiles of the sequencing libraries were examined by Agilent Bioanalyzer 2100 using a High Sensitivity DNA chip (Agilent). Two indexed libraries were equimolarly pooled and sequenced on Illumina NOVAseq 6000 platform using the v2 Kit (Illumina, San Diego, CA) with a customized paired end, dual indexing (26/8/0/98-bp) format according the manufacturer’s protocol. Using proper cluster density, a coverage around 250 M reads per sample (2000–5000 cells) were obtained corresponding to at least 50.000 reads per cell.

## Statistical analysis

Statistical analyses were performed using PRISM (GraphPad, version 6.0). Statistical significance was tested with the unpaired (nonparametric) t test. N, p-values, and significance are reported in each figure and legend. All results were expressed as means ± SD.

## Data analysis

Single cell sequenced libraries were aligned with CellRanger pipeline, we used Hg38 for indexing reference transcripts. Resulting data was imported in python as anndata object. We used scanpy vs 1.3.1 and pandas for downstream analyses in python. Normalization was performed following Seurat^48^ pipeline. From the resulting data we removed hematopoietic cells selected for: 1) clustering together using UMAP algorithm 2) expressing high levels of PTPRC (figS8a). For visualization purposes, neighbors were calculated with 900 n_neighbors and plotted with UMAP and diffusion map.

For graphs in Figure 4d we calculated z score considering all the cells regardless of the condition. In the left panels we kept the diffusion coordinates and considered the same color scale for all the graph. The same z score was used in the frequency plot on the right, plotted data was normalized dividing by the area under the curve.

Single cells data was than clustered by diffusion distance to obtain more coverage and more consistent results for the downstream analyses. Final number of cells per cluster is between 20 and 40, these numbers were chosen to minimize the differences in number of reads per cluster and in order not to lose the heterogeneity of the system. Data was imported in R and Gene groups were identified with WGCNA (with minClusterSize 42 and SoftPower 5) considering among the Module-Trait Relationships (MTRs) those with high correlation between patients or conditions.

Differential expression was performed with edgeR, filtering genes by FDR < 0.05 and logFC > 1.25, to keep all the possible information for the downstream analysis (gene ontologies, pathways analysis, etc).

We used a combination of Causal Network Analysis, Downstream Effects Analysis, Upstream Regulator Analysis and Molecule Activity Predictor from Ingenuity Pathway Analysis^42^ to identify the impact of the genes identified from differential expression or from correlation network analysis.

Heatmaps were generated from logCPM using a modified version of Clustergrammer^49^, hierarchical clusterings, were included, were performed considering correlation as distance.

## Code availability

All code used for analysis is available from the corresponding author upon reasonable request.

## Data availability

All RNAseq data presented in this study will be made available in the recommended public repositories upon publication.

## Supporting information

## Acknowledgements

This work was supported by the Associazione Italiana per la Ricerca sul Cancro (AIRC) (IG 2014-2018 to G.T. and E.C.V., IG-14622 to U.C.); EPIGEN Flagship Project of the Italian National Research Council (CNR) (to G.T., E.C.V., R.L., M.D.S., R.J.P.B., S.M., S.P. and M.P.); the European Research Council (DISEASEAVATARS to G.T.); Fondazione Umberto Veronesi (to T.V.); the Fondazione IEO-CCM; Centro Cardiologico Monzino to M.L.; Fondazione Italiana per la Ricerca sul Cancro (FIRC) (to P.L.R.); the Fondazione Cariplo (grant 2017-0886 to A.L.T.); AIRC grant n° IG2016-ID18575 and ERC Consolidator Grant n° 617978 to M.P. the Italian Ministry of Health (Ricerca Corrente grant to G.T., Ricerca Finalizzata PE-2016-02362551 to U.C.). We are grateful to IEO Biobank staff for collection of surgical samples. We are grateful to Stefano Freddi for technical support (flow cytometry). We are grateful to Cristina Cheroni and Alessandro Vitriolo for discussion on data analytical tools.

## Author Contributions

T.V. performed the experiments. C.E.V. developed and implemented the computational methods and conducted the analysis for scRNAseq. M.L. and U.C. provided background knowledge on HGSOC stem cells, M.L. helped with setting conditions for 2D cultures; P.L.R contributed to the experimental and analytical design and provided expertise on the biological model; R.L. contributed to discussion on the experimental set-up; A.L.T. and M.D.S. provided expertise and help for the preparation of scRNAseq libraries for the first sample. S.P.,M.P. and S.M. provided intellectual input in setting up the Organoid-focused EPIGEN Flagship Project within which this study unfolded; R.B. assisted in the computational setup. N.C. headed the HGSOC surgery and supervised the provision of clinical informations. T.V., U.C. and G.T. developed the experimental design for the monoclonal 3D culture; T.V., C.E.V. and G.T. designed the experiments, critically evaluated the results and wrote the manuscript; G.T. conceived and supervised the study.

## Declaration of interests

The authors declare no competing interests.

## Supplementary table S1

**Table S1.** Table listing markers for: cancer initiating cells (CICs), epithelial mesenchymal transition (EMT) and epithelial cells (EPI). CICs: CD44 antigen (CD44), aldehyde dehydrogenase family member A1 (ALDH1A1), prominin-1 or CD133 antigen (CD133), sex determining region Y-box 2 (SOX2), ecto-5′ - nucleotidase or CD73 (NT5E), homeobox protein NANOG (NANOG), notch homolog 1, translocation-associated (Drosophila) (NANOG), tyrosine-protein kinase receptor 1 (ROR1). EMT: twist-related protein 1 (TWIST), snail family transcription repressor 1(SNAI1) snail family transcription repressor 2 (SNAI2), zinc finger E-box binding domain homeobox 1 (ZEB1), vimentin (VIM), cadherin 2 (CDH2), integrin subunit beta 6 (ITGB6), fibronectin 1 (FN1), zinc finger E-box binding domain homeobox 2 (ZEB 2). EPI: keratin 18 (KRT18), cadherin 1 (CDH1), mucin 1 (MUC1), cancer antigen 125 (CA125),epithelial cell adhesion molecule (EPCAM), mesothelin (MSLN), keratin 7 (KR7), keratin 8 (KR8).

| CICs | EMT | EPI |
| --- | --- | --- |
| CD44<sup>47</sup> | TWIST<sup>48</sup> | KRT18<sup>6</sup> |
| ALDH1A1<sup>49</sup> | SNAI1<sup>50</sup> | CDH1<sup>51</sup> |
| CD133<sup>49</sup> | SNAI2<sup>50</sup> | MUC1<sup>52</sup> |
| SOX2<sup>53</sup> | ZEB1<sup>54</sup> | CA125<sup>52</sup> |
| NT5E<sup>18</sup> | VIM<sup>55</sup> | EPCAM<sup>52</sup> |
| NANOG<sup>56</sup> | CDH2<sup>55</sup> | MSLN<sup>57</sup> |
| NOTCH1<sup>47</sup> | ITGB6<sup>55</sup> | KRT7 |
| ROR1<sup>57</sup> | FN1<sup>55</sup> | KRT8 |
|  | ZEB2<sup>58</sup> |  |

**Table 1.** Clinical-pathological parameters of patients. Overview of the diagnosis, the disease state and the grade of the tumor for each patient with HGSOC from which scOCOCs were generated. Age of the patient is referred at the time of diagnosis.

| Sample | Diagnosis | Disease state | Grade | Age |
| --- | --- | --- | --- | --- |
| Patient 1 | Serous surface papillary carcinoma | Primary | 3 | 60 |
| Patient 2 | Serous surface papillary carcinoma | Primary | 3 | 46 |
| Patient 3 | Serous cystadenocarcinoma | Primary | 3 | 56 |
| Patient 4 | Serous surface papillary carcinoma | Primary | 3 | 62 |
| Patient 5 | Serous cystadenocarcinoma | Primary | 3 | 58 |

## Supplementary figure

Correlation of the percentage of the ascitic fluid and the growth of scOCOs. Briefly, single cell derived from 2D cultures from three different patients with HGSOC were plated in 96 well plates in presence of STEM media only (0) and stem media supplemented with different volume of ascitic fluid 10, 25, 50, 100 and 200 □l (only ascites). We observed that the best working solution to reach the higher number of scOCOs is obtained adding a volume between 25/50 □l of ascitic fluid. All the experiments were performed supplementing 25 ml of ascitic fluid up to a final volume of 200 □l per well. Results represent the mean ± SD of number of wells with scOCOs of three independent experiments performed on three different samples.

Supplementary figure 2. Ascitic fluid supplemented to 2D primary culture increase cell proliferation. (a) Bright field image of 2D cultured cells plated with culturing media for tumor primary cells supplemented with ascitic fluid (left picture), 2D primary culture plated in their growth media only (right panel), x2 magnification. (b) Cell proliferation was correlated with cell metabolic activity measured by MTT assay. Cells derived from HGSOC ascites were plated in 96 well collagenated plates at different number in their culture media supplemented (purple bars) or not (pink bars) with ascitic fluid. Cell proliferation was measured 24 hours following administration by MTT. Results represents the mean ± SD of two experiments performed on two different samples in triplicate.

Supplementary figure 3: Bar plots showing single cell organoid forming efficiency (scOFE) in three different patients of two scOCOs derived from different patients at passage 1 and passage 2. One different organoid at passage 1 P1 was used to generate two different scOCOs at passage 2 (P2 and P2*). Results represent the mean ± SD of three idependent experiments performed on three different samples.

Supplementary figure 4. Ascitic fluid (AS) from different patients do not affect single cell organoids forming efficiency. Graph bar showing scOFE as percentage, briefly single cell derived from 2D cultures from 2 different patients with HGSOC were plated in three different 96 well plates, one in presence of the ascitc fluid derived from the same patient (AS from the same patients), the second and the third with the ascitic fluid from other two unrelated patients with HGSOC (AS from unrelated patient 1-2) scOFE as percentage was calculated dividing the number of the wells with HGSOC for the number of the total cells plated. Results represent the mean ± SD of two separate experiments performed on two different samples.

Supplementary figure 5: (a) Enrichment analysis of epithelial signatures. Left panel: diffusion map of all the categories and patients with a color scale defined by the z score of the epithelial signature; right panel: frequency plot showing the variation in the distribution of cells as a function of the z score of the indicated signature. (b) Percentage of cells with expression z score for the epithelial signature among categories (yellow: fresh ascites, turquoise: 2D primary cells, purple: single cell derived ovarian cancer organoids or scOCOs) at different passages (P1 panel on the top right and P2 panel on the bottom right).

Supplementary figure 6: Patient Specificity.(a) Venn diagram of differentially expressed genes, in at least one of the patients, shared between the indicated conditions.(b) WGCNA labeled heatmap of whole transcriptome. Color maps define correlations between groups of genes and patients. (c) UMAP of all the conditions calculated using the aggregation of gene modules in fig. S6b blue, green and greenyellow. Color code indicates patients. (d) UMAP of all the conditions calculated using the aggregation of gene modules in fig. S6b blue, green and greenyellow. Color code indicates culture conditions.

**uFig. 1.**
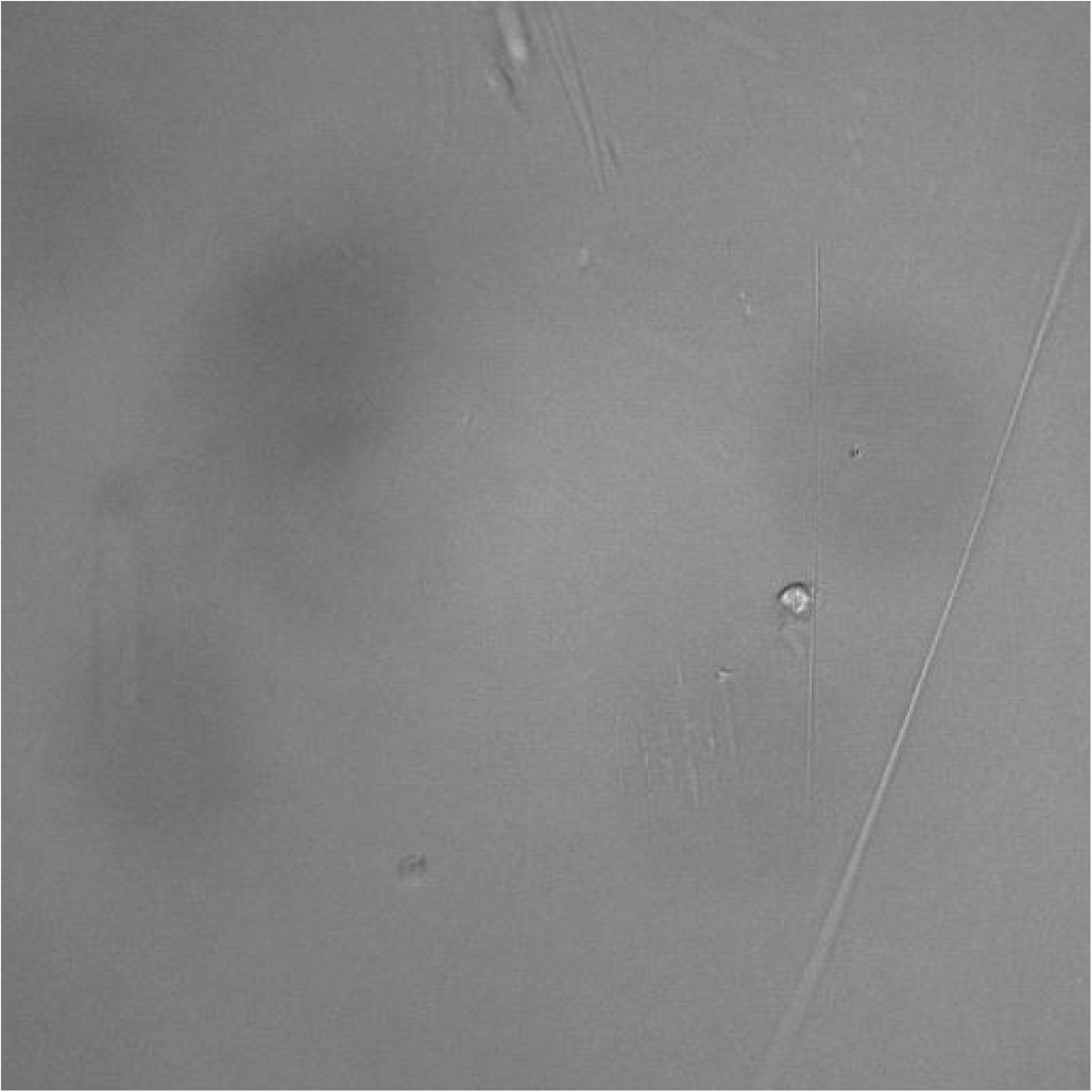
Box initially at rest on sled sliding across ice.

