## Supplementary material for "Single cell derived organoids capture the self-renewing subpopulations of metastatic ovarian cancer"

### Supplementary figure 1

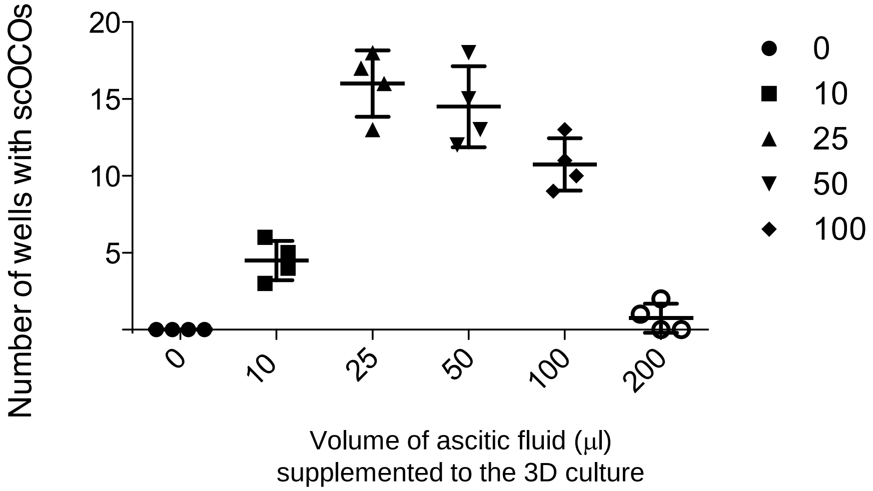

Supplementary figure 2

**a**

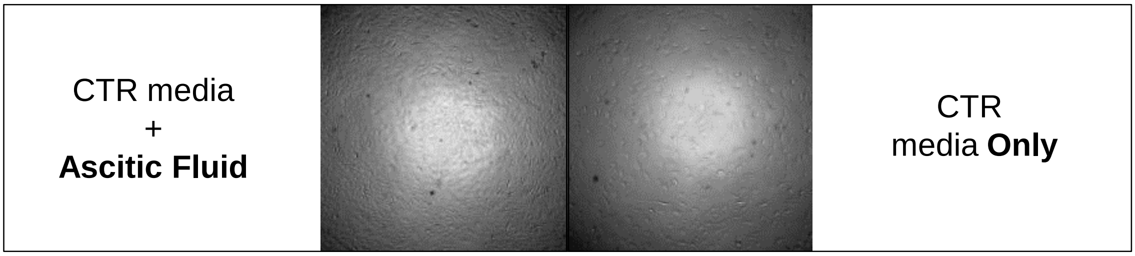


**b**

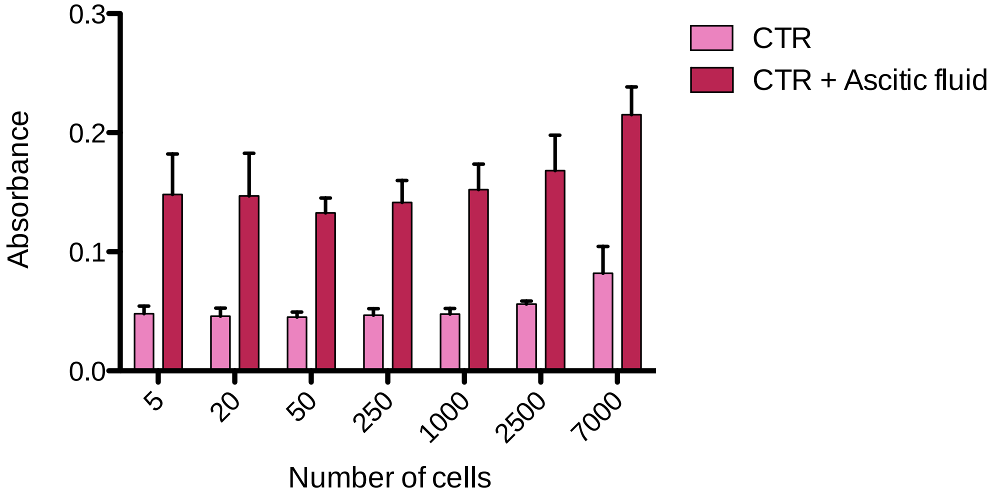

### Supplementary figure 3

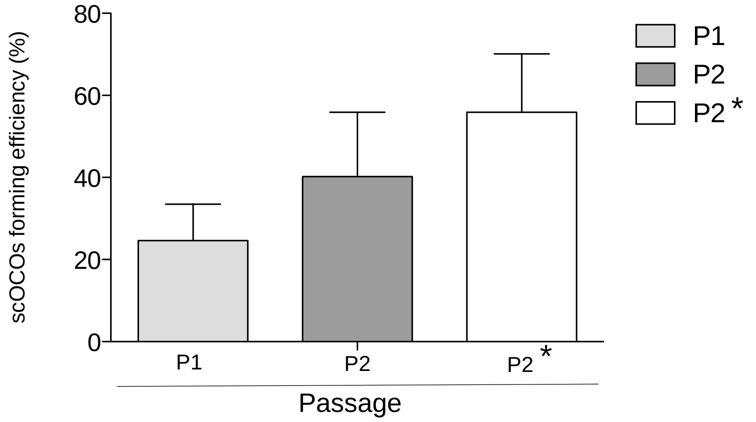

### Supplementary figure 4

#### Ascitic fluid from different patients do not affect single cell organoid forming efficiency

(2D-scOCOs from patient 5)

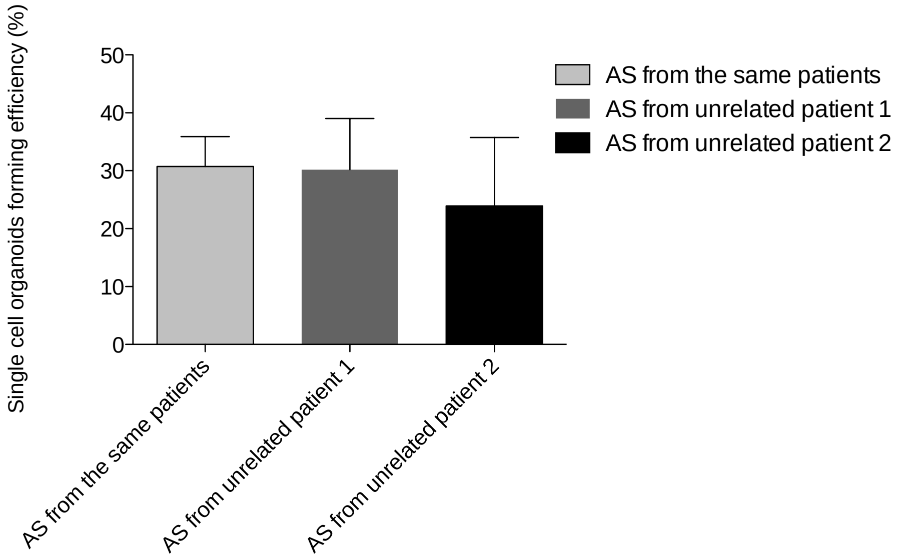

Supplementary figure 5

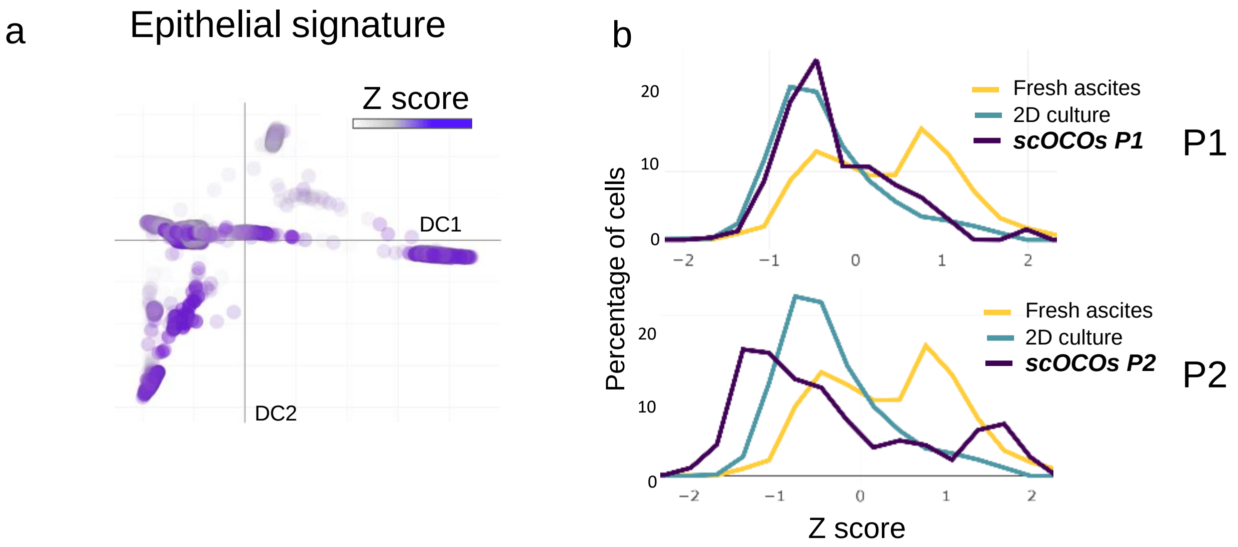

### Supplementary figure 6

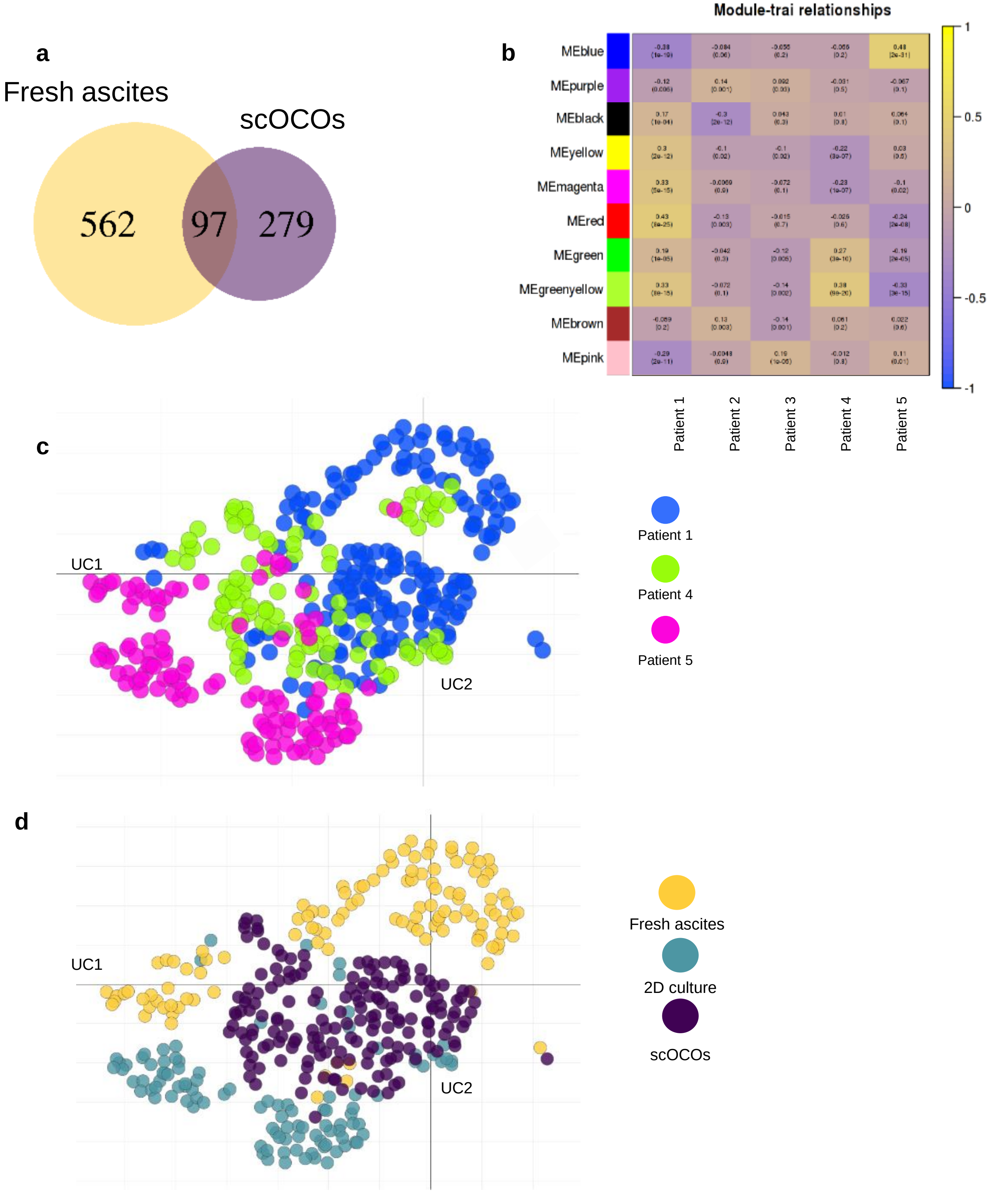

### Supplementary figure 7

**a**

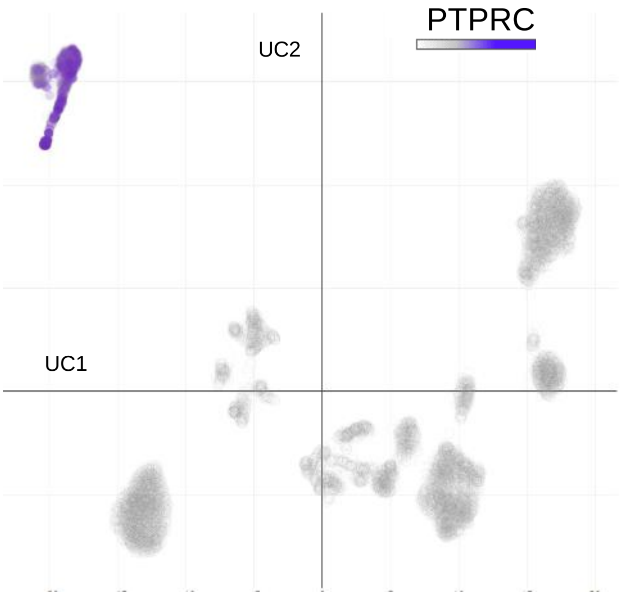
